# Characterization of FGFR signaling in prostate cancer stem cells and inhibition via TKI treatment

**DOI:** 10.1101/2020.12.13.422586

**Authors:** J. Ko, A. N. Meyer, M. Haas, D. J. Donoghue

## Abstract

**Background:** Metastatic castrate-resistant prostate cancer (CRPC) remains uncurable and novel therapies are needed to better treat patients. Aberrant Fibroblast Growth Factor Receptor (FGFR) signaling has been implicated in advanced prostate cancer (PCa), and FGFR1 is suggested to be a promising therapeutic target along with current androgen deprivation therapy.

**Methods:** We established a novel *in vitro* 3D culture system to study endogenous FGFR signaling in a rare subpopulation of prostate cancer stem cells (CSCs) in the cell lines PC3, DU145, LNCaP, and the induced pluripotent iPS87 cell line.

**Results:** 3D-propagation of PCa cells generated spheroids with increased stemness markers ALDH7A1 and OCT4, while inhibition of FGFR signaling by BGJ398 or Dovitinib decreased cell survival and proliferation of 3D spheroids. The 3D spheroids exhibited altered expression of EMT markers associated with metastasis such as E-cadherin, vimentin and Snail, compared to 2D monolayer cells. TKI treatment did not result in significant changes of EMT markers, however, specific inhibition of FGFR signaling by BGJ398 showed more favorable molecular-level changes than treatment with the multi-RTK inhibitor Dovitinib.

**Conclusions:** This study provides evidence for the first time that FGFR1 plays an essential role in the proliferation of PCa CSCs at a molecular and cellular level, and suggests that TKI targeting of FGFR signaling may be a promising strategy for AR-independent CRPC.

## INTRODUCTION

Prostate cancer (PCa) is the fifth leading cause of cancer-related deaths among men worldwide ^1^. PCa is considered to be a hormone sensitive disease, with androgen receptors as the central therapeutic target. Androgen deprivation therapy (ADT) is the first-line of treatment for both non-metastatic and metastatic PCa patients. Apalutamide, a potent AR inhibitor recently approved by the U.S. Food and Drug Administration, demonstrates significant clinical efficacy, increasing metastasis-free patient survival by 24.3 months ^2^. However, while ADT is effective at initial stages, androgen-independent tumor cells eventually emerge and most patients relapse and develop metastatic castrate-resistant prostate cancer (mCRPC) ^3^.

Among tumor cells, a rare subpopulation of cells exhibiting stem/progenitor properties are believed to be responsible for cancer recurrence, metastasis and chemo-resistance ^4-7^. These rare populations of cells are known as cancer stem cells (CSCs) or tumor-initiating cells for their ability to generate tumors *in vivo* with high efficiency. Cancer stem cells have been characterized with respect to biomarker expression, such as cell surface markers, functional markers such as self-renewal genes, and intracellular enzyme activity, which may be responsible for drug resistance ^8^. Cells that are propagated in three-dimensional (3D) culture possess the ability to grow in an anchorage-independent manner; these cells also exhibit increased stem and progenitor-like properties and the ability to undergo epithelial-to-mesenchymal transition (EMT) ^7-9^. ALDH7A1 is a commonly used biomarker to identify CSCs in PCa ^10^.

3D spheroid cultures select for CSCs and exhibit advantages over conventional 2D cell culture and animal models. Compared to 2D culture, the tumor spheroids and organoids can better recapitulate the natural structure and heterogeneity of a tumor, which may contains different stages of proliferating cells and a necrotic core with chemical gradients of oxygen and nutrients. 3D spheroid cultures also exhibit clinically more relevant prediction in drug testing ^5,11-13^.

Fibroblast Growth Factor Receptors (FGFRs) are members of the receptor tyrosine kinase (RTK) family and consist of FGFR1, 2, 3 and 4, encoded by four different genes. Binding of FGF ligands along with heparin sulfate proteoglycans to the receptors triggers their dimerization and trans-autophosphorylation. In turn, this initiates downstream signal transduction cascade including activation of PLCγ, PI3K/AKT, RAS/MAPK, and JAK/STAT pathways. These pathways regulate many biological responses, such as embryonic development, cell proliferation, differentiation, survival, mitogenesis and angiogenesis ^14,15^. However, aberrant FGFR activation has been implicated in numerous developmental diseases and various cancers including prostate, breast, ovarian, gastric cancer and glioblastoma presenting FGFR inhibition an attractive therapeutic target ^14,16,17^. In PCa, loss of PTEN, a tumor suppressor gene, and overactivation of Akt are frequently observed, which is suggested to be responsible for chemotherapy and radiation resistance and tumor invasion and metastasis ^18,19^.

FGFR signaling has been associated with promoting stem cell-like properties in various cancers such as breast cancer ^20,21^, non-small cell lung cancer ^22^, and esophageal squamous cell carcinoma ^23^. However, despite some important studies, the importance of FGFR signaling in prostate CSCs remains unclear. Prior research has reported that FGFR1 is upregulated in CRPC patient samples and is associated with higher relapse rates and poor survival ^24^. Others have reported that FGFR1 and FGFR4 were overexpressed in PCa patient samples and showed that inhibition of FGFR4 decreased cell proliferation and invasion in a DU145 cell line study ^25^. Another study, using mouse models, suggested that FGFR1 activation drives PCa progression and EMT ^26^. Lastly, it was shown that reported that FGFR1 was upregulated in clinical prostate tumor samples, and treatment with tyrosine kinase inhibitors (TKIs) showed promising antitumor effects depending on FGFR1 expression ^27^.

In this study, we introduce a novel 3D culture model to investigate whether FGFR signaling is required for cell survival and proliferation of prostate CSCs. We have examined 3D spheroids of common PCa cell lines, PC3, DU145 and LNCaP, and spheroids of patient-derived iPS87 cells, a novel induced pluripotent stem (iPS) cell line ^28,29^. Using unique suspension culture conditions without the ectopic addition of growth factors for culturing 3D spheroids, we evaluated the effects of TKIs PD166866, BGJ398, and Dovitinib. The findings provided in this study provide a better understanding of the importance of FGFR signaling in PCa.

## RESULTS

### FGFR Expression and Downstream Signaling in Spheroids of PC3, LNCaP and DU145 cells

PC3, LNCaP and DU145 cells are the most commonly used PCa cell lines and are derived from bone, lymph node, and brain metastases, respectively. PC3 and DU145 are highly metastatic and AR-negative, whereas LNCaP is less tumorigenic and AR-positive ^30^. Due to the heterogeneous nature of PCa, we set out to investigate and characterize all three cells lines.

First, we examined the expression of each FGFR as several studies have reported that overexpression of FGFR1 and FGFR4 were observed and associated with PCa progression and metastasis ^24-27^. In this study utilizing cell lines, only DU145 cells showed significant expression of FGFR1 in 2D monolayer culture (Figure 1A, 1^st^ row panels). We also examined the expression of other FGFR family members and detected only FGFR4 (Figure 1A, 4^th^ row panels). This finding was consistent with prior data showing that FGFR4 is predominantly expressed in PC3, DU145 and LNCaP cell lines, and not FGFR1, with the exception of DU145 cells ^31^.

**Figure 1.**
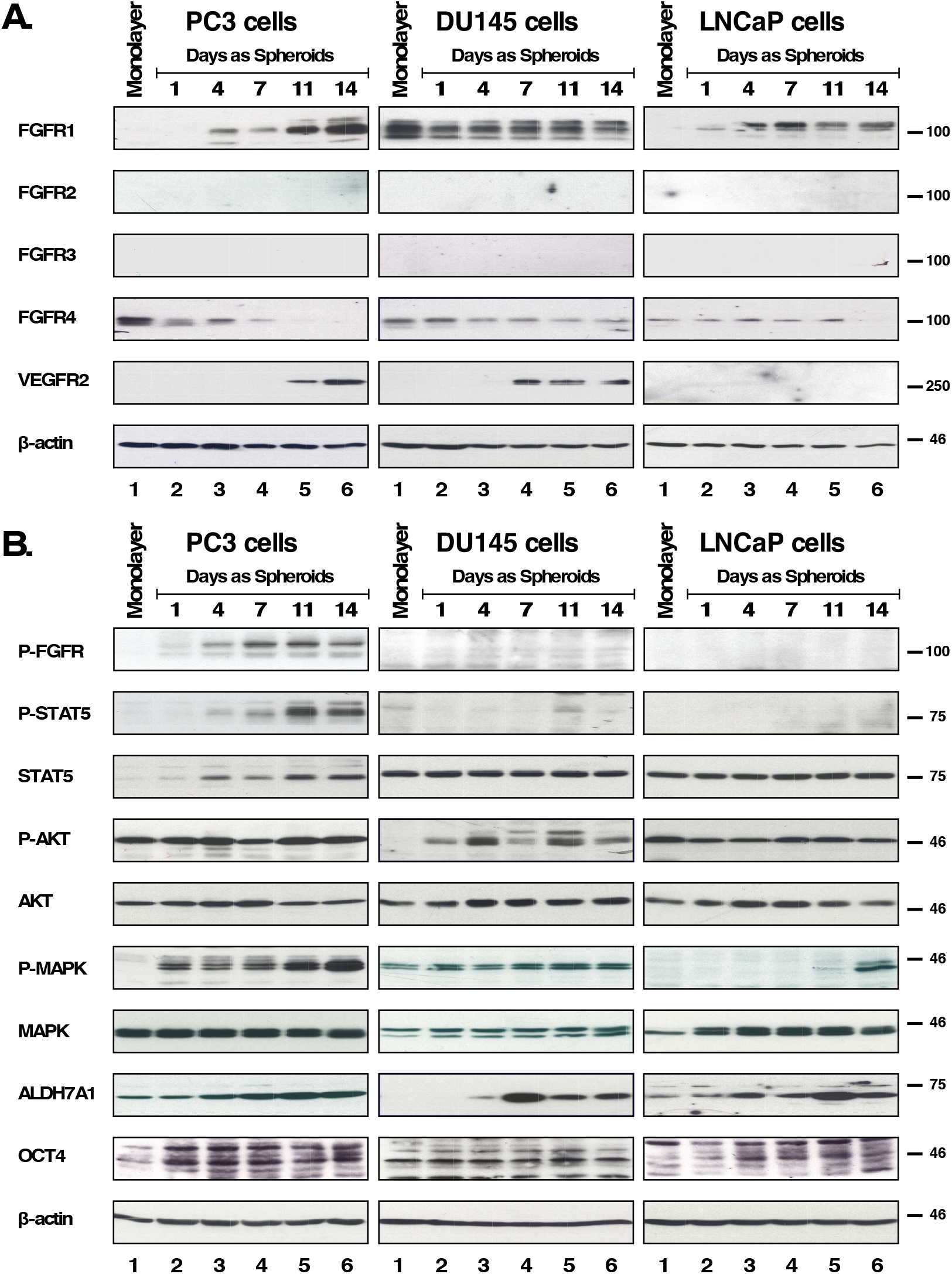
Expression and downstream cell signaling activation of FGFR of spheroids of PC3, DU145 and LNCaP. **(A)** 2D monolayer cells and 3D spheroids of PC3, DU145 and LNCaP cells on days 1, 4, 7, 11, and 14 were subjected to Westernblot analysis for FGFR 1-4, and VEGFR2. Beta-actin was used as a loading control. M= 2D monolayer. **(B)** (1st row) FGFR activation was shown by immunoblotting for phospho-Y653/654 FGFR antiserum and only PC3 spheroids showed positive signal. (2nd row) STAT5 activation was detected by immunoblotting for phospho-Y694-STAT5, and the same membrane was stripped and probed for total STAT5 expression shown immediately below. (4th row) AKT activation was detected by immunoblotting for phospho-S473-AKT, and the same membrane was stripped and probed for total AKT expression shown immediately below. (6th row) MAPK activation was shown by immunoblotting for phospho-T202/Y204-MAPK, and the same membrane was stripped and probed for total MAPK shown immediately below. (8th row) ALDH7A1 expression. (9th row) OCT4 expression. (10th row) Beta-actin was used as a loading control.

We utilized sphere formation assays to select for a rare subpopulation of prostate CSCs, to determine whether FGFR protein expression is different compared to typical monolayer-cultured 2D cells. Abundant evidence suggests that 2D monolayer cells display altered gene and protein expression, and possess different properties of proliferation, angiogenesis, cell interaction, and drug sensitivity compared to 3D-culture cells ^12,32,33^. We utilized agarose gel-coated tissue culture dishes to select for anchorage-independent growth using RPMI 1640 medium supplemented with 10% serum replacement, commonly used in culturing embryonic stem cells and induced pluripotent stem cells ^34^. The spheroids were collected on days 1, 4, 7, 11, and 14 and were subjected to immunoblotting to observe changes in protein expression.

We probed for protein expression of FGFR1-4 and, additionally, Vascular Endothelial Growth Factor Receptor 2 (VEGFR2), another RTK in the VEGFR family. VEGFR2 was of interest due to its association with increased malignancy, and its function as a regulator of angiogenesis, invasion, metastasis, development, and bone destruction ^35^. Furthermore, VEGFR2 is also one of the main targets of Dovitinib, one of the TKIs examined in this study. As determined by immunoblotting, we found that PC3 spheroids exhibited increasing FGFR1 and VEGFR2 expression while FGFR4 expression decreased with increasing days in culture (Figure1A, left panels). DU145 spheroids maintained both FGFR1 and FGFR4 expression and showed increasing VEGFR2 expression (Figure 1A, middle panels). LNCaP spheroids also showed increasing FGFR1 but no detectable expression of VEGFR2 (Figure 1A, right panels). Of note, spheroids of AR-negative PCa cell lines, PC3 and DU145, either increased or maintained FGFR1 expression, respectively, and both exhibited increased VEGFR2 expression. In all three cell lines, FGFR2 and FGFR3 were not detected. Beta-actin was used as a loading control (Figure 1A).

Next, we examined activation of signaling pathways by immunoblotting with p-FGFR antisera and downstream effectors using p-STAT5, p-MAPK, and p-AKT antisera. We found FGFR kinase activity was most significant in PC3 spheroids, exhibiting upregulated p-FGFR, and p-STAT5, and p-MAPK (Figure 1B. Left panels). LNCaP spheroids showed activation of the MAPK pathway, while DU145 spheroids showed activation of the AKT pathway, neither of which was activated in monolayer cells. In PCa cells AKT has been shown to negatively regulate MAPK ^36^, consistent with our findings for the monolayer cells that when p-AKT is observed, p-MAPK is not, and vice versa. Unexpectedly, however, we detected activation of both pathways in spheroids (Figure 1B. Middle and right panels).

As the reliability of cell surface markers remains debatable in PCa CSCs, we utilized functional markers such as ALDH7A1 and OCT4 instead of cell surface markers ^37-39^. ALDH7A1 detoxifies aldehyde compounds induced by chemotherapeutic agents to be associated with CSC markers in PCa, and is highly expressed in primary tumors and the matched bone metastases of those primary tumors ^40^. We observed that spheroids of all 3 cell lines exhibited an increase in ALDH7A1 (Figure 1B, 8^th^ panels). OCT4, an embryonic stem cell marker for self-renewal and maintenance of an undifferentiated state, was maintained throughout the 3D culture with little increase (Figure 1B, 9^th^ panels). Beta-actin was used as a loading control (Figure 1B, 10^th^ panels).

Taken together, the different protein expression profiles between 2D monolayer cells and 3D spheroids cells support the idea that anchorage-independence confers a unique advantage in examining the importance of FGFR signaling in PCa CSCs.

### Effect of FGFR Inhibition on Proliferation of Spheroids

Due to FGFR activation seen in Figure 1B, we examined the requirement of FGFR signaling for cell survival and proliferation of PC3, DU145, and LNCaP spheroids. PC3, DU145, and LNCaP spheroids were treated with TKIs Dovitinib, BGJ398, or PD166866. Both Dovitinib and BGJ398 have been used in clinical trials for several cancers with defined FGFR genetic alterations, while PD166866 is a highly selective inhibitor towards FGFR1 and other kinases such as c-Src, PDGFR, and EGFR ^41^.

PC3, LNCaP, and DU145 monolayer cells were propagated as spheroids and Dovitinib, BGJ398 or PD166866 were added to the cultures at low, medium and high concentrations every 3-4 days. Proliferation of the cultures was determined using the metabolic MTT assay, described in Material and Methods. At 14 days of culture, spheroids were counted and analyzed from random field views by bright field microscopy.

Figure 2A shows the different cell morphology between the PC3 monolayer cells and PC3 spheroids at 14 days. PC3 spheroids exhibit grape-like or loose clusters of cells indicating poor cell-cell contact. Dovitinib (0.5-2 µM) demonstrated potent anti-proliferative effects as shown in Figure 2B. Interestingly, PC3 spheroids displayed a differing morphology with granular nuclei and larger cell size when treated with 2µM Dovitinib (data not shown). PC3 cells treated with BGJ398 (1-5µM) also showed inhibitory effects (Figure 2C); although not as effective as the other two inhibitors, while PC3 cells treated with PD166866 (2-20 µM) successfully hindered cell proliferation at a much higher concentration (Figure 2D).

**Figure 2.**
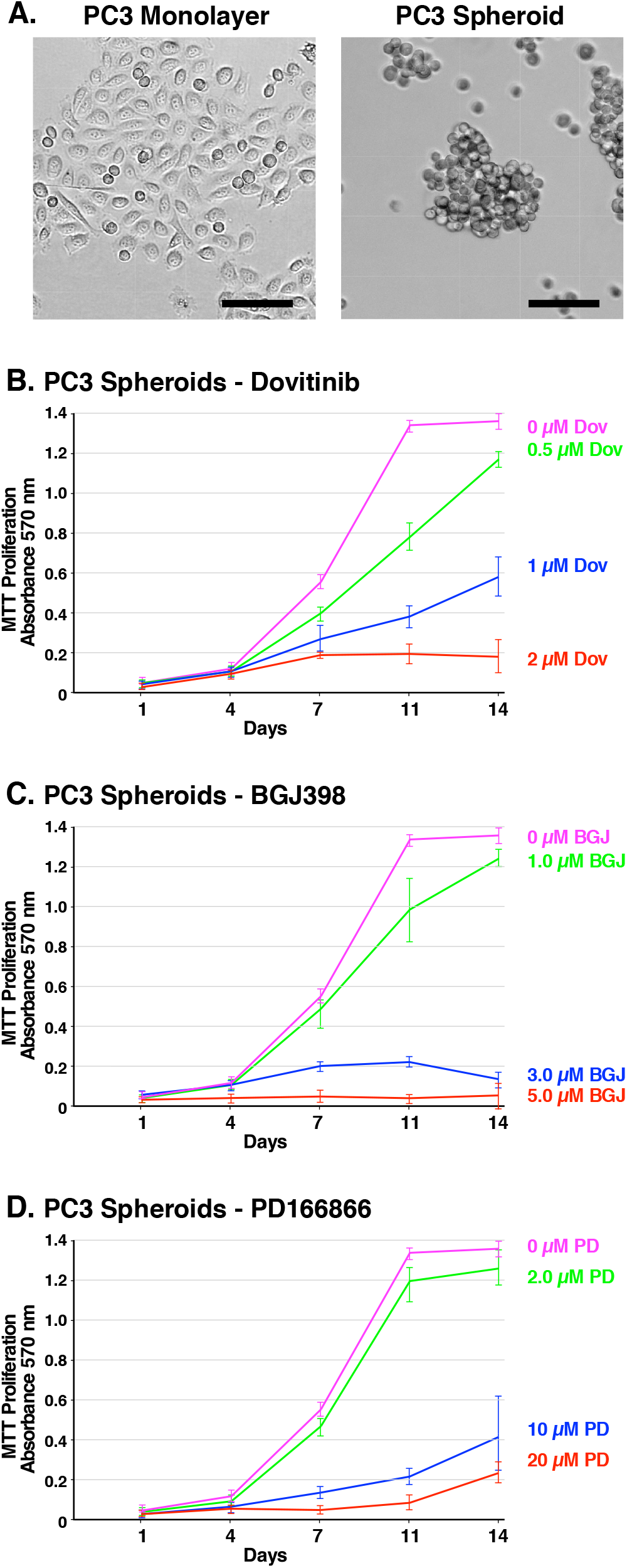
Formation of PC3 spheroids and inhibition of survival and growth via TKI treatment. **(A)** Brightfield microscope images of PC3 cells. Left; 2D monolayer. Right; 3D spheroids at 14 days. The scale bars indicate 100 µm. **(B-D)** Biological triplicate cultures of PC3 3D spheroids were grown in RPMI 1640 with 10% SR on agarose-coated dishes. Samples of cultures were taken and assayed by MTT metabolic assay indicating the number of viable cells on days 1, 4, 7, 11, and 14 to show the proliferation over time. **(B)** 2μM -20μM of PD166866 was treated. **(C)** 1μM -5μM of BGJ398 was treated. **(D)** 0.5μM -2μM of Dovitinib was treated. Error bars show the standard deviation. PD= PD166866, BGJ = BGJ398, Dov = Dovitinib.

Figure 3A shows the different cell morphology between the DU145 monolayer cells and DU145 spheroids at 14 days. DU145 spheroids are tightly packed together indicating robust cell-to-cell adhesion. Furthermore, they have an irregular, round shape and are smaller in size than LNCaP spheroids. DU145 cells exhibited a noticeably lower capacity for anchorage-independent growth than PC3 and LNCaP cells. Furthermore, DU145 spheroids were sensitive to treatment with Dovitinib, BGJ398, and PD166866, suggesting that FGFR activation is required for their proliferation (Figure 3B-D).

**Figure 3.**
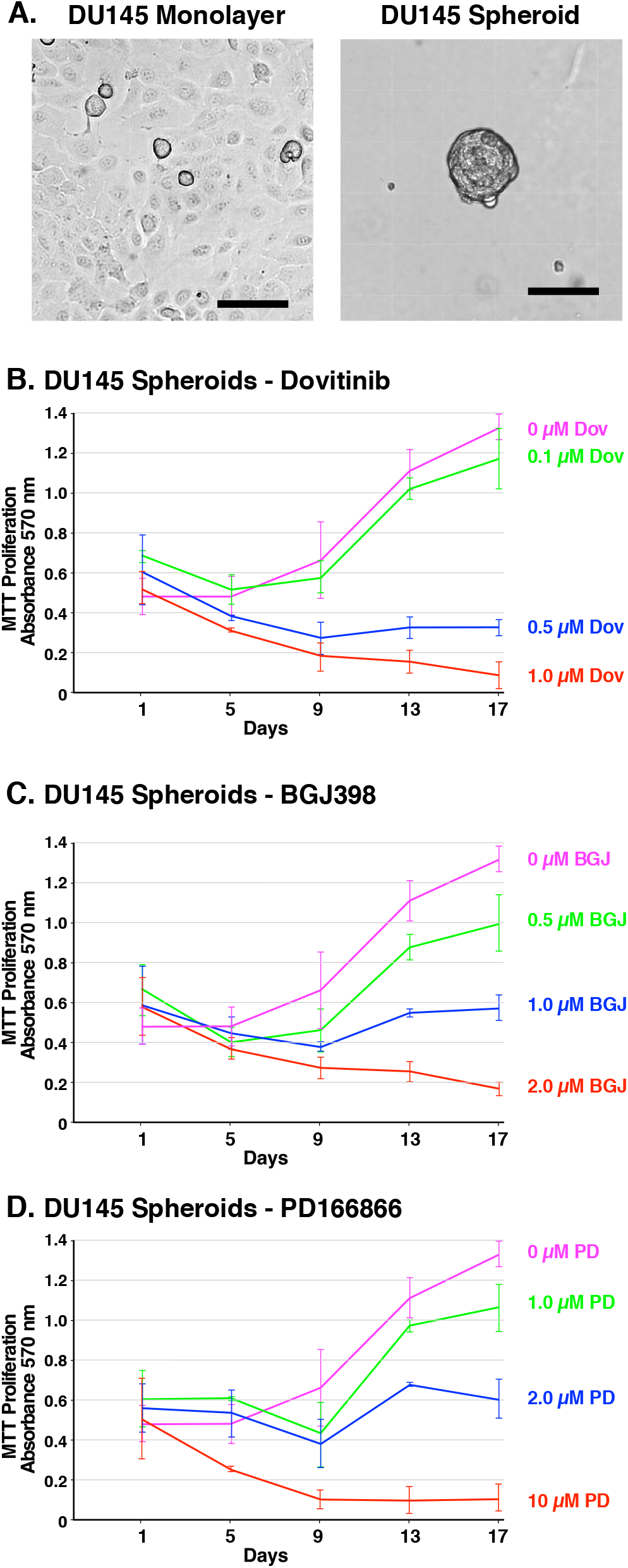
Formation of DU145 spheroids and inhibition of survival and growth via TKI treatment. **(A)** Brightfield microscope images of DU145 cells. Left; 2D monolayer. Right; 3D spheroids at 14 days. The scale bars indicate 100 µm. **(B-D)** Biological triplicate cultures of PC3 3D spheroids were grown in RPMI 1640 with 10% SR on agarose-coated dishes. Samples of cultures were taken and assayed by MTT metabolic assay indicating the number of viable cells on days 1, 4, 7, 11, and 14 to show proliferation over time. **(B)** PC3 spheroids were treated with 2μM - 20μM PD166866. **(C)** PC3 spheroids were treated with 1μM - 5μM BGJ398. **(D)** PC3 spheroids were treated with 0.5μM - 2μM Dovitinib. Error bars show the standard deviation. PD= PD166866, BGJ = BGJ398, Dov = Dovitinib.

Figure 4A shows the cell morphology of LNCaP monolayer cells and LNCaP spheroids at 14 days. LNCaP spheroids exhibit robust cell-to-cell adhesion, forming round-shaped masses. LNCaP cells are characterized by slow proliferation, and required a much higher seeding density compared to PC3 cells. Anti-proliferative effects were observed in response to Dovitinib (0.5-2µM). However, LNCaP spheroids were less responsive to treatment with BGJ398 (2.5-10µM). The inhibitory effect of PD166866 (2-20µM) was similar to PC3 spheroids (Figure 4B-D). Taken together, these data demonstrate the inhibitory potential of Dovitinib, BGJ398, and PD166866 through an anti-proliferative mechanism, particularly for AR-negative cell types.

**Figure 4.**
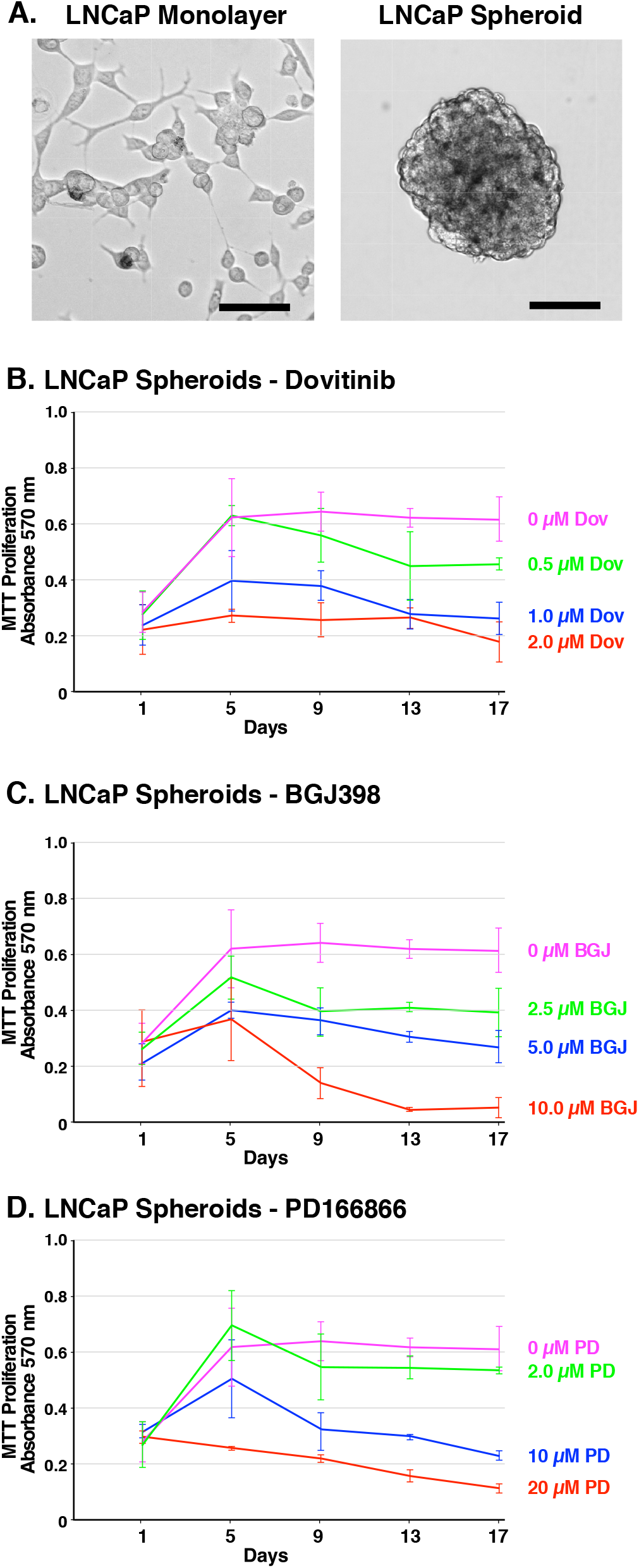
Formation of LNCaP spheroids and inhibition of survival and growth via TKI treatment. **(A)** Brightfield microscope images of LNCaP cells. Left; 2D monolayer. Right; 3D spheroids at 14 days. The scale bars indicate 100 µm. **(B-D)** Biological triplicate cultures of LNCaP 3D spheroids were grown in RPMI 1640 with 10% SR on agarose-coated dishes. As described in Material and Methods, samples of cultures were taken and assayed by MTT metabolic assay indicating the number of viable cells on days 1, 5, 9, 13, and 17 to show the proliferation over time. **(B)** LNCaP spheroids were treated with 2μM - 20μM PD166866. **(C)** LNCaP spheroids were treated with 2.5μM - 10μM BGJ398. **(D)** LNCaP spheroids were treated with 0.5μM -2 μM Dovitinib. Error bars show the standard deviation. PD= PD166866, BGJ= BGJ398, Dov= Dovitinib.

### Effect of FGFR Inhibition on Signaling Pathways of Spheroids

Immunoblotting was used to access the activation of FGFR signaling pathways in inhibitor-treated PC3, DU145, and LNCaP spheroids. PC3, DU145, and LNCaP spheroids were treated with either Dovitinib, PD166866, or BGJ398 and collected after 14 days. Spheroids were then subjected to immunoblotting.

In PC3 spheroids, we found that PD166866 did not effectively suppress the p-FGFR signal compared to the control, showing its poor efficacy even at a high concentration of 10µM (Figure 5A. Panel 3, lanes 2-4). FGFR phosphorylation was most successfully suppressed with BGJ398, an FGFR-selective inhibitor, followed by Dovitinib treatment, as seen by immunoblotting (Figure 5A. Panel 3, lanes 6, 8).

**Figure 5.**
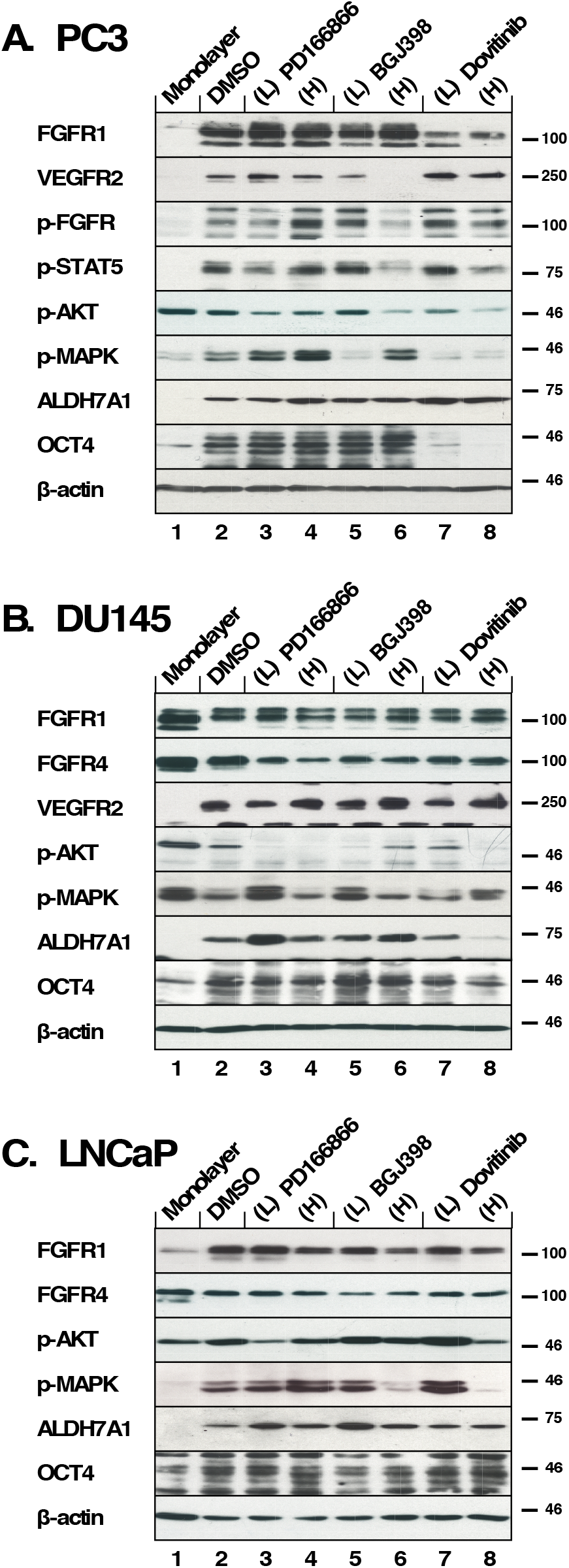
The effects of FGFR TKI treatment on 3D spheroid signaling pathways. All inhibitors were added every 3-4 days during 2 weeks of culture in biological triplicate and lysates were subjected to western blot analysis. **(A)** PC3 spheroids were treated with PD166866 was at (L) 2 µM, (H) 10 µM; with BGJ398 at (L) 1 µM, (H) 3 µM; or with Dovitinib at (L) 0.5 µM, (H) 1 µM concentrations. (Panels 1, 2) Expression of FGFR1 and VEGFR2; (Panel 3) p-FGFR signal was detected using p-Y653/654 antibody; (Panel 4) p-STAT5 detected by p-Y694 antibody; (Panel 5) p-AKT was probed using p-S473 antibody; (Panel 6) p-MAPK signal shown by phospho-p44/42 MAPK (Erk1/2) (T202/Y204); (Panel 7) Expression of ALDH7A1; (Panel 8) Expression of OCT4 expression; (Panel 9) Beta-actin as a loading control. **(B)** DU145 spheroids were treated with PD166866 at (L) 1 µM, (H) 2 µM; with BGJ398 at (L) 0.5 µM, (H) 1 µM; or with Dovitinib at (L) 0.1 µM, (H) 0.3 µM concentrations. (Panels 1, 2, and 3) Expression of FGFR1 FGFR4 and VEGFR2; (Panel 4) p-AKT; (Panel 5) p-MAPK; (Panel 6) Expression of ALDH7A1; (Panel 7) OCT4 expression; (Panel 8) Beta-actin as a loading control. **(C)** LNCaP spheroids were treated with PD166866 at (L) 2 µM, (H) 8 µM; with BGJ398 at (L) 2.5 µM, (H) 5 µM; or with Dovitinib at (L) 0.5 µM, (H) 1 µM concentrations. (Panels 1, 2) Expression of FGFR1 and FGFR4; (Panel 3) p-AKT; (Panel 4) p-MAPK signal; (Panels 5, 6) Expression of ALDH7A1 and OCT4; (Panel 7) Beta-actin as a loading control.

Both p-STAT5 and p-AKT were reduced by the high concentrations of BGJ398 and Dovitinib (Figure 5A. Panels 4 & 5, lanes 6, 8). Interestingly, the p-MAPK signal increased with the high concentration treatment of BGJ398, perhaps suggesting a compensatory mechanism in PC3 spheroids allowing survival (Figure 5A. Panels 6, lanes 5-8).

ALDH7A1 expression was largely not affected by FGFR inhibition except for a slight increase with Dovitinib treatment (Figure 5A. Panel 7, lanes 2-8). As shown in Figure 1B, PC3 spheroids exhibit increased OCT4 expression compared to the 2D-cultured cells and, interestingly, an increase in OCT4 expression is seen with Dovitinib treatment (Figure 5A. Panel 8, lanes 2-8). We suggest that Dovitinib treatment may have an adverse effect of inducing PC3 spheroids into a more aggressive neuroendocrine phenotype via non-FGF receptor signaling ^42^.

When DU145 spheroids were examined for their responses to TKI treatment, FGFR phosphorylation was not detected. We examined the expressions of FGFR1, FGFR4, and VEGFR2 and found no noticeable changes (Figure 5B. Panels 1-3, lanes 2-8). As DU145 spheroids were sensitive to all three TKIs, we speculate that FGFR signaling is critical in supporting the survival and growth of these spheroids; however, FGFR total expression may be below our detection limit by immunoblotting. We observed little activation of AKT (Figure 5B. Panel 5, lanes 2-8), and irregular activation of MAPK signaling in DU145 spheroids (Figure 5B. Panel 6, lanes 2-8).

ALDH7A1 expression in DU145 spheroids was largely unaffected by FGFR inhibition, with the exception of Dovitinib treatment. The decrease in ALDH7A1 expression (Figure 5B. Panels 6, lanes 2-8) may indicate a reduction of proliferative potential. However, OCT4 expression was maintained regardless of TKI treatment, suggesting that properties of stemness may not be altered by TKI treatment (Figure 5B. Panel 7, lanes 2-8).

We also examined the responses to the FGFR TKIs using AR-positive LNCaP spheroids. We observed no change in expression of FGFR1 and 4 (Figure 5C. Panels 1, 2, lanes 2-8). As a result of FGFR inhibition, AKT activation was largely unaffected (Figure 5C. Panel 3, lanes 2-8), however, MAPK activation was ablated (Figure 5C. Panel 4, lanes 6, 8). These data suggest a mechanism by which LNCaP spheroid proliferation is inhibited by suppressed MAPK activation, although AKT activation persists correlating with their survival.

LNCaP spheroids did not exhibit noticeable changes in ALDH7A1 or OCT4 expression in the presence of high TKI concentrations (Figure 5C. Panels 5, 6, lanes 2-8); this suggests that the AR-positive LNCaP spheroids may not depend on FGFR signaling as much as AR-negative prostate cancer cells.

Taken together, we found that the CSC markers ALDH7A1 and OCT4 were up-or downregulated with Dovitinib treatment differently between PC3 spheroids and DU145 spheroids, while the FGFR-selective inhibitor BGJ398 did not show a disparity between these cell lines. This result may be explained by non-specific effects of Dovitinib on other RTKS than FGFRs; this may underlie the differences observed in ALDH7A1 and OCT4 expression in the different cell types. The identification of these differences may help screening for a molecular subgroup of PCa that would be predictive of different treatment outcome by Dovitinib.

### Effect of FGFR Inhibition on Gene Expression of 3D spheroids of AR-independent Prostate Cancer Cell Lines

We were interested in investigating whether AR-independent PC3 and DU145 spheroids would exhibit altered transcriptional responses to TKI treatment. Target genes examined by RT-qPCR included FGFR1, OCT4, ALDH7A1, E-cadherin, Vimentin, and Snail. Relative mRNA expression was determined by normalizing against the housekeeping gene, beta-actin (Figure 6. A-F).

**Figure 6.**
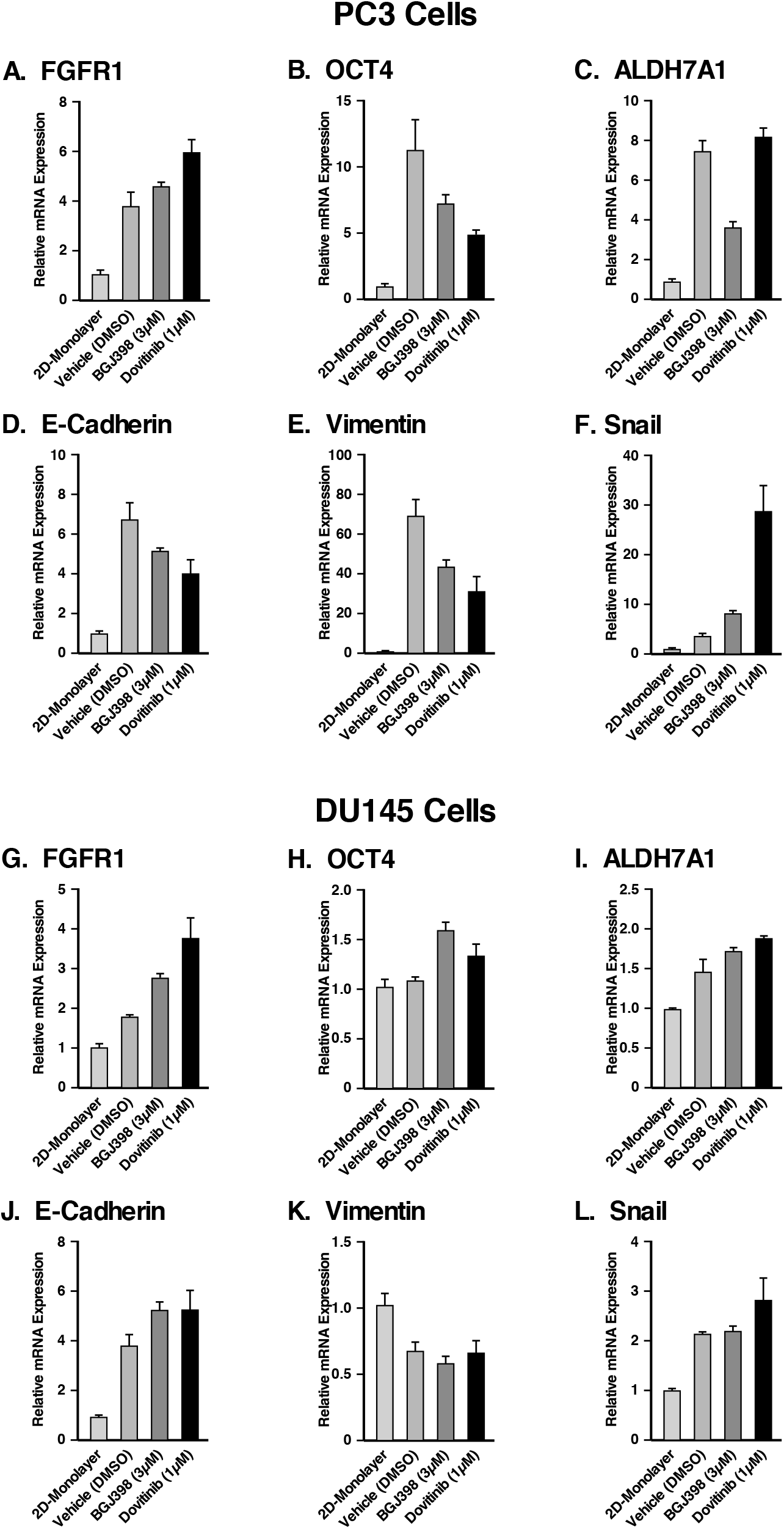
Gene expression of the spheroids of PC3 and DU145 cell lines compared to 2D monolayer culture and their response to FGFR inhibition. RT-qPCR was performed on PC3 cells grown in a monolayer, and PC3 cells grown as spheroids either with or without TKI treatment. The following target genes were analyzed through mRNA expression: **(A)** FGFR1, **(B)** OCT4, a stem cell marker, **(C)** ALDH7A1, prostate specific CSC marker, and **(D)** the epithelial marker, E-cadherin, **(E, F)** the mesenchymal markers, vimentin and Snail. mRNA expression was normalized against beta-actin and the fold changes were evaluated. RT-qPCR was performed on DU145 cells grown in a monolayer, and DU145 cells grown as spheroids either with or without TKI treatment. The following target genes were analyzed through mRNA expression: **(G)** FGFR1, **(H)** OCT4, **(I)** ALDH7A1, **(J)** E-cadherin, **(K)** vimentin and **(L)** Snail. mRNA expression was normalized against beta-actin and the fold changes were evaluated. The data represent the average of three biological independent experiments and the error bars represent standard error of the means (mean ± SEM).

In PC3 spheroids (Figure 6. A-F) FGFR1 mRNA was increased about 3.8-fold in PC3 spheroids compared to 2D monolayer cells, suggesting that FGFR1 supports the anchorage-independent proliferation of PC3 spheroids. Treatment with BGJ398 and Dovitinib resulted in FGFR1 up-regulation, possibly compensating for TKI inhibition. Propagation of PC3 cells as spheroids in comparison with 2D-cultured cells exhibited up-regulation of OCT4 about 11-fold, and ALDH7A1 about 7-fold, showing that PC3 spheroids proliferation upregulates genes that promote stemness. Treatment with BGJ398 and Dovitinib resulted in decreased OCT4, suggesting that FGFR1 regulates stemness in these spheroids. ALDH7A1 mRNA levels were decreased with BGJ398 treatment, however, ALDH7A1 mRNA levels remained the same with Dovitinib treatment. Taken together, inhibition of FGFR1 via TKI treatment appears to target the CSC population of PC3 spheroids, suppressing the self-renewal ability of these cells.

Using RT-qPCR, the mRNA levels of EMT markers E-cadherin, Vimentin, and Snail were examined in 3D spheroids and 2D-cultured cells. Results showed significant up-regulation of E-cadherin, vimentin, and Snail in PC3 spheroids. Notably, PC3 spheroids showed about 69-fold increase in the level of vimentin ^43^. BGJ398 and Dovitinib treatment reduced the mRNA levels of E-cadherin and vimentin, but not of Snail. Notably, Dovitinib treatment increased the expression of Snail significantly compared to the negative control. Taken together, these results indicate that the specific inhibition of FGFR signaling by BGJ398 showed more favorable molecular-level changes than treatment with the multi-RTK inhibitor Dovitinib, in PC3 spheroids.

FGFR1 mRNA levels increased by nearly 2-fold in DU145 spheroids when compared to 2D cultured cells. Additionally, FGFR1 mRNA levels increased further with TKI treatment (Figure 6. H-M). Little change was observed in OCT4 expression, while ALDH7A1 mRNA level increased about 1.5-fold. These results indicate that TKI treatment had little effect on genes correlated with stemness in DU145 spheroids.

Similar to PC3 spheroids, DU145 spheroids up-regulated E-cadherin, and Snail in comparison with 2D-cultured cells; however, the level of vimentin decreased about 0.6-fold. No significant changes were observed in the EMT markers in response to TKI treatment. Although it was observed that DU145 spheroids respond to TKI treatment as assessed by proliferation, TKI treatment did not lead to any significant changes in the gene regulation shown by the RT-qPCR results.

In conclusion, higher fold changes were observed for CSC and EMT markers in PC3 spheroids than in DU145 spheroids compared to their 2D monolayer cells, and the effects of TKI treatment were more prominent in PC3 spheroids. Nonetheless, these data support the conclusion that CSCs enriched through spheroid proliferation exhibit properties that correlate with metastatic potential, in comparison with 2D-cultured cells.

### Utilization of Induced Pluripotent Stem (iPS) 87 Cells

As we examined enriched CSC populations from the common cancer cell lines PC3, DU145, and LNCaP by 3D spheroid culture, we concurrently investigated the iPS87 cell line which originated from a PCa biopsy sample ^28,29^. This cell line was established by reprogramming primary prostate tumor cells into induced pluripotent stem cells and was shown to possess tumor initiating ability *in vivo* ^29^. iPS87 cells were propagated as previously described ^29^. Single iPS87 cells detached from the mouse embryonic feeder layer after one day, and by 7 days these cells were able to propagate into spheroids (Figure 7A). By 14 days they were fully grown, and no apparent size change was observed at later times.

**Figure 7.**
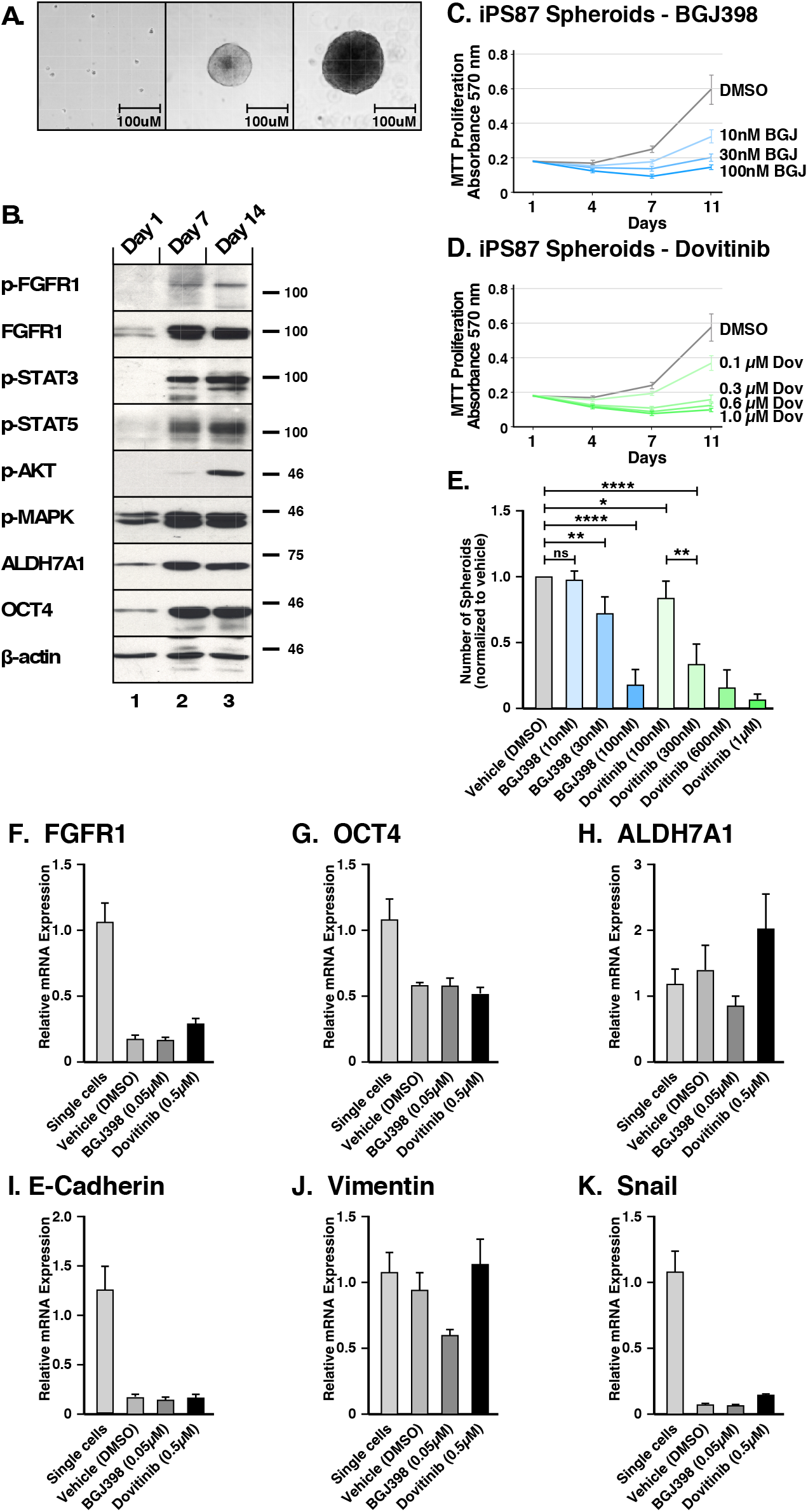
FGFR signaling in induced Pluripotent Stem (iPS) 87 Spheroids. **(A)** Left panel shows single cells of iPS87 after 24 hours of culturing on a TC plate in ES+/+ medium; middle panel shows growth of a spheroid at day 7; right panel shows a spheroid at day 14. The scale bars indicate 100 µm. **(B)** iPS87 cells on day 1 as single cells in lane 1, 3D spheroids on day 7 in lane 2, and 3D spheroids on day 14 in lane 3 were immunoblotted for: (1st row) FGFR activation was shown by immunoblotting for phospho-Y653/654 FGFR antiserum; (2nd row) Total FGFR1 expression was is shown; (3rd row) STAT3 activation was detected by immunoblotting for phospho-Y705-STAT3; (4th row) STAT5 activation was detected by immunoblotting for phospho-Y694-STAT5; (5th row) AKT activation was detected by immunoblotting for phospho-S473-AKT; (6th row) MAPK activation was shown by immunoblotting for phospho-T202/Y204-MAPK; (7th row) Total ALDH7A1 expression; (8th row) OCT4 expression is shown; (9th row) Beta-actin was used as a loading control. **(C-E)** Triplicate cultures of 3D spheroids of iPS87 were grown in ES+/+ on 12-well TC plates. Samples of cultures were assayed by MTT metabolic assay indicating the number of viable cells on days 1, 4, 7, and 11 to show the proliferation over time. **(C)** iPS87 spheroids were treated with 10nM - 100nM of BGJ398. **(D)** iPS87 spheroids were treated with 100nM - 1μM of Dovitinib. All experiments were performed in biological triplicate; error bars show standard deviation. **(E)** The number of spheroids (>1,000 µm^2^) from each well of 12-well plates was determined by counting at day 11. Error bars show standard deviation. *P* values are from two-tailed paired t tests. ns= not significant (*P*>0.05), * = *P* ≤0.05, ** = *P* ≤0.01, **** = *P* ≤0.0001. **(F-L)** RT-qPCR was performed on iPS87 cells grown as single cells, and iPS87 cells grown as spheroids either with or without TKI treatment. The following target genes were analyzed through mRNA expression: **(F)** FGFR1, **(G)** OCT4, **(H)** ALDH7A1, **(I)** E-cadherin, **(J)** vimentin, and **(K)** Snail. mRNA expression was normalized against beta-actin and the fold changes were evaluated. Data represent average of three independent experiments; error bars represent standard error of the means (mean ± SEM).

The protein expression of iPS87 single cells and spheroids was examined on days 1, 7, and 14 (Figure 7B). It was found that the iPS87 cells initially express FGFR1, but activated p-FGFR was not detectable (Panels 1 and 2). Interestingly, iPS87 spheroids display an increase in FGFR1 expression, and a phospho-FGFR signal was readily detected by immunoblotting. Additionally, activation of STAT3, STAT5, AKT, and MAPK pathways was also detected (Panels 3-6). Similarly, an increase of expression in CSC markers ALDH7A1 and OCT4 was also detected in in iPS87 spheroids (Panels 7 and 8).

To examine the effect of BGJ398 and Dovitinib, MTT metabolic assay was conducted on days 1, 4, 7 and 11 after propagation in the presence of the TKIs. Both BGJ398 (10nM-100nM) and Dovitinib (100nM-1μM) demonstrated an inhibitory effect, reducing proliferation in a dose-dependent manner (Figure 7C, D). iPS87 spheroids responded to TKI treatment at nanomolar range, highlighting their dependency of FGFR signaling. The number of spheroids also decreased as a result of TKI treatment (Figure 7E). Collectively, these data demonstrate that FGFR inhibition results in decreased cell proliferation of iPS87 cells.

### Effect of FGFR Inhibition on Gene Expression of 3D spheroids of iPS87 cells

In order to quantify gene expression, RT-qPCR was performed on monolayer-cultured cells and 3D spheroids of iPS87 cells, in the absence or presence of the TKIs BGJ398 and Dovitinib. RT-qPCR was performed to detect FGFR1, OCT4, ALDH7A1, E-cadherin, Vimentin, and Snail. The relative mRNA expression was determined by normalizing against the housekeeping gene, beta-actin (Figure 7F-K).

Noticeably, FGFR1 mRNA decreased about 0.18-fold in spheroids compared to single cells; this was in contrast to the observed upregulation of FGFR1 protein expression demonstrated by immunoblotting. The spheroids also showed a decrease in OCT4 mRNA (∼0.59-fold), in contrast to up-regulation of OCT4 protein, as seen by immunoblotting. However, TKI treatment of iPS87 did not affect mRNA levels of FGFR1 or OCT4. The level of ALDH7A1 mRNA showed no significant changes as the iPS87 cells grow into spheroids or when treated with TKIs.

We examined the mRNA levels of EMT markers of E-cadherin, vimentin, and Snail similarly as for PC3, DU145, and LNCaP cell lines. The results showed that E-cadherin and Snail were significantly down-regulated in the spheroids but were not affected by TKI treatment. Vimentin level was not changed by spheroid propagation and was only affected when treated with BGJ398, being decreased by about 0.59-fold. Overall, FGFR inhibition did not result in significant changes in EMT markers; thus, although FGFR inhibition reduced the survival and proliferation of the spheroids, no dramatic molecular-level changes were observed which would correlate with the suppression of FGFR signaling and inhibition of cellular proliferation observed in response to TKI treatment.

## DISCUSSION

### Prostate Cancer as a Stem Cell Disease

Current PCa therapies focus on targeting AR signaling. However, a majority of patients eventually develop progressive disease, eventually becoming AR-negative and castration independent. For these patients, ADT is no longer a plausible treatment option. This highlights the need for the development of novel therapeutic options for Pca. The heterogeneity of PCa, in contrast to many other solid tumors, presents many challenges. Efforts have been made to find non-AR related targets that can be clinically utilized for prognostic purposes, and FGFR1 has been shown to be a promising alternative target ^24-27^.

It is generally believed that PCa tumors and cell lines contain a rare population of CSCs, which are AR-independent and exhibit properties of stemness ^28,44^. While ADT may be effective against more differentiated AR-positive cells, the reoccurrence of cancer may be due to this subpopulation of AR-independent CSCs. In this study, we wished to investigate the possible importance of FGFR signaling in the proliferation and maintenance of PCa CSCs. Additionally, the study aimed to compare different FGFR inhibitors, differing in their specificity and off-target effects. For these studies, we employed a 3D culture system that promotes the proliferation of spheroids using several commonly studied PCa cell lines, together with a patient-derived pluripotent stem cell line iPS87 ^29^.

### FGFR Signaling in 3D-cultured Spheroids

Through the implementation of 3D culture conditions, the results presented here demonstrate for the first time that FGFR1 expression is up-regulated in PC3 and LNCaP spheroids and is maintained in DU145 spheroids. This establishes a new 3D *in vitro* model to study the involvement of FGFR1 using common PCa cell lines. Although FGFR1 is overexpressed in PCa patient samples, as seen through sequencing and FGFR1 PDX models ^27^, typical PCa cell lines do not exhibit detectable FGFR1. Furthermore, the TKI AZ8010 has previously been shown to inhibit the growth of PCa cells expressing FGFR1 or FGFR4, suggesting that targeting FGFR signaling inhibits PCa progression ^31^. Despite these advances, the *in vitro* model presented here utilizing 3D spheroids may offer significant advantages to aid in the investigation of the importance of FGFR in PCa.

The 3D culture of spheroids provides the opportunity to select for CSCs, providing a cell population enriched for stem and progenitor like cells in comparison with 2D-cultured monolayers, as seen through the data presented here. This study demonstrates the importance of FGFR signaling in PCa CSCs that require FGFR signaling for their survival and proliferation. Both FGFR specific inhibitors and multi-RTK inhibitors effectively suppressed the proliferation of spheroids in the AR-negative PC3 and DU145 cell lines, and in the pluripotent iPS87 cell line. This work further suggests that AR-negative prostate cancers may depend on FGFR for oncogenic cell growth. Previously, FGFR1 was shown to promote stemness in malignant subpopulations in lung cancer ^45^ and breast cancer ^21^. Results presented here demonstrate for the first time that FGFR1 supports the CSC population in PCa not only at a cellular level but also at a molecular level.

Dovitinib has been shown to target FGFR1 in PCa *in vivo*, however the effects of Dovitinib on FGFR downstream signaling pathways such as STAT, AKT, and MAPK are not understood ^27^. Results presented here demonstrate up-regulation of endogenous FGFR1 in CSC-enriched spheroids, facilitating the study of FGFR inhibition. Using PCa cell lines, PC3, Du145, and LNCaP, we show that 3D-cultured spheroids exhibited different levels of activation of the STAT, AKT, and MAPK pathways. TKI treatment with either BGJ398 or Dovitinib consistently decreased downstream signaling activation.

Of note, LNCaP spheroids failed to respond significantly to TKI treatment, despite the up-regulation of FGFR1, which may be due to the inefficient inhibition of AKT signaling shown by immunoblotting. Moreover, as LNCaP cells are AR-positive, they may activate compensatory signaling mechanisms. Prior work has shown that AR-null/NE-null LNCaP cells which survived AR antagonists exhibited increased FGF signaling and were more sensitive to FGFR inhibition than AR-positive LNCaP cells ^46^. Furthermore, MAPK pathway activation was identified as the principal mechanism supporting proliferation and survival of the double negative PCa cells *in vitro* and *in vivo*. Taken together, results from prior work together with those presented here strongly suggest that FGFR/MAPK inhibition may be a promising strategy for the treatment of AR-negative PCa, but less effective for AR-positive PCa.

Furthermore, the differential activation of FGFR and its downstream signaling between 2D and 3D cell cultures provides evidence that FGFR signaling is involved in maintenance of PCa stem cells. Numerous studies have emphasized that cells grown as 2D-monolayers may not recapitulate gene and protein expression and thus serve as an inaccurate representation of drug response in human tumor physiology ^12,13^. This study demonstrates PC3, DU145 and iPS87 3D spheroids show differential activation of FGFR as well as up-regulation of OCT4, a stem cell marker, and ALDH7A1, a CSC marker ^10,40^, when examined by RT-qPCR.

As CSCs are considered to be responsible for drug resistance in many cancer types including PCa, this study aimed to examine whether TKI treatment targeting FGFR would show signs of drug resistance suggested via ALDH7A1 expression. This study demonstrated that Dovitinib treatment increased both gene and protein expression of ALDH7A1 in PC3, DU145 and iPS87 spheroids. These results suggests that Dovitinib treatment may have adverse effects and that the use of a more specific TKI against FGFR may be a more favorable approach. This may be related to the ability of Dovitinib to promote neuroendocrine differentiation in PC3 and LNCaP cells, representing a more aggressive phenotype ^42^. Interestingly, treatment with BGJ398, an FGFR selective inhibitor, showed a decrease in FGFR expression in PC3 and iPS87 spheroids. However, further investigation is needed to assess direct correlation of cancer stemness in relation to TKI treatment.

### EMT Markers

The results show that PC3 spheroids significantly up-regulate Vimentin and Snail, critical mesenchymal markers in the EMT process, suggesting increased metastatic potential of cells that are enriched for stemness by 3D-propagation as spheroids. Our results are consistent with prior work, describing re-expression of previously downregulated E-cadherin in advanced PCa, which allows E-cadherin to serve as a potential biomarker of disease progression ^47^. Additional studies have shown that E-cadherin positive subpopulations in PC3 and DU145 cell lines, exhibited highly invasive properties, significant tumor formation in mouse models, and higher expression of stem cell markers ^48^. Taken together, results reported here reinforce the utility of 3D-propagation to produce spheroids cell cultures enriched in PCa CSCs ^9^.

DU145 spheroids showed an increase in E-cadherin and Snail, similar to PC3 spheroids; however, the levels of vimentin decreased. This may simply reflect the heterogeneity of PCa cell lines and human PCa in general. Further examination of mesenchymal markers such as ZEB, Twist, and N-cadherin may provide a more complete understanding of EMT and metastasis for this cell line. As for iPS87 cells, the gene expression of single cells of iPS87 which possess stemness and self-renewal properties was compared to that of 3D spheroids consisting of necrotic core, proliferating zone, and the outer layer, which requires cellular differentiation. Although the 3D spheroids of iPS87 may appear to have decreased stemness properties compared to single cells, based on the reduced gene expression of OCT4 and ALDH7A1, the protein expression of these markers was significantly elevated. Therefore, despite the decreased gene expression of E-cadherin and Snail, further examination of these markers would provide a more complete understanding of the proliferative potential of the spheroids.

Collectively, FGFR inhibition via BGJ398 and Dovitinib did not induce any significant changes of EMT marker expression of the 3D spheroids of PC3, Du145, and iPS87. Additional *in vivo* assays which assess the effects of FGFR inhibition on gene expression will be valuable to further understand the role of TKIs with respect to EMT in PCa.

Previous studies have addressed the lack of patient-specific *in vitro* models that accurately reflect the diversity of human PCa,, which has hampered the development of effective treatment. Thus, utilizing patient-derived organoids (PDOs) and iPS-derived organoids (iDOs) have emerged as promising strategies for disease modeling and drug development. In a prior study from our group, iPS-derived cells were obtained from human PCa biopsy samples to produce iPS87 cells ^29^; this may provide a useful platform for other researchers to study PCa stem cells with restored tumor initiating properties *in vivo*. It was hypothesized that iPS87 spheroids may be utilized as iDOs for their characteristics of self-organizing free-floating cell aggregates with higher order tissue complexity. The term organoid has not been clearly defined with regard to PCa until the recent successful generation of a fully mature organoid that appears to recapitulate human prostate; this study showed that human iPSC-derived cells that are co-cultured with rodent urogenital sinus mesenchyme by 12 weeks can comprehensively generate prostate tissue with epithelial architecture, including cells at different differentiation stages of basal and luminal cells as well as neuroendocrine cells ^49^.

In conclusion, based on our studies of 3D-cultured spheroids propagated from PC3, DU145, LNCaP, and iPS87 PCa cells, we suggest that TKI targeting of FGFR signaling may be a promising strategy for AR-independent CRPC. The study provides evidence for the first time that FGFR1 plays a role in supporting the proliferation of PCa CSCs at a molecular and cellular level.

## MATERIALS AND METHODS

### Cell Lines and Culture

DU145 and LNCaP cells were obtained from American Type Culture Collection and PC3 were obtained from Dr. Leonard Deftos at UCSD, and maintained in RPMI 1640 media supplemented with 10% fetal bovine serum and 1x pen/strep and in 5% CO_2_ at 37°C. For spheroid assays, cells were maintained in RPMI 1640 media supplemented with 10% Gibco KnockOut Serum Replacement (KnockOut SR) from ThermoFisher Scientific.

Induced Pluripotent Stem (iPS) 87 cells (iPS87) were generated as previously described ^28,29^. The iPS87 cells were grown on Mitomycin-C inactivated MEF feeder cells and maintained in KnockOut DMEM (Gibco) supplemented with 0.125% Bovine Serum Albumin (Sigma), 2% L-glutamine, 1% non-essential amino acids, 1% Fungizone/0.5% gentamycin 10% serum replacement (Gibco), 6.25 ng/mL bFGF (Peprotech), referred to as “ES+/+,” 5% CO2, 37°C ^29^.

### Spheroid Assays

Single cells of PC3, DU145 and LNCaP were obtained by dissociation with Versene/EDTA incubation and seeded onto 1% agarose-coated TC plates in 10% SR/RPMI 1640 media with 1X pen/strep with the following densities: PC3 at an initial density of ∼6.6 x 10^3^ cells/ml, DU145 at ∼3.3x 10^4^ cells/ml, and LNCaP at ∼3.3 x 10^4^ cells/ml. The iPS87 spheroids were cultured in ES+/+ media on tissue culture dishes without MEF feeder cells to readily grow into spheroids from single cells at density of 8.0 x 10^4^ cells/ml. Images of spheroids (and monolayer cells) were acquired using an inverted microscope (Carl Zeiss Microscopy GmbH, Germany) with a 20x objective. Image processing was done in Fiji/ImageJ.

### MTT Metabolic Assays and Addition of Inhibitors

From the initial plating of single cells onto the non-adherent substrates, measurements were taken after 1, 4, 7, 11, and 14 days for PC3 cells or after 1, 5, 9, 13, and 17 days for DU145 and LNCaP cells. PC3 cells were plated on 60 mm plates with total 3 ml volume of media, at each time point 300 μl of cell cultures were transferred to a 24-well TC plate with an additional 200 μl of media and incubated with 50 μl of 5 mg/mL of thiazolyl blue tetrazolium bromide (MTT) (Sigma) at 37°C in 5% CO_2_ for 4 h, after which 500 μl of 0.04 M HCl in isopropanol was added and incubated again for at least 30 min. Absorbance was measured at 570 nm. DU145 and LNCaP cells were plated on 10 cm plates with a total volume of 10 ml of media, at each time point 1 ml of cultures were transferred to a 24-well TC plate and incubated with 100 μl of 5 mg/mL of thiazolyl blue tetrazolium bromide (MTT) at 37°C in 5% CO2 for 4 h, and cells were concentrated by centrifugation and removing 770 μl of the supernatant, after which 300 μl of 0.04 M HCl in isopropanol was added and incubated again for at least 30 min. Absorbance was measured at 570 nm. Inhibitors (PD166866, BGJ398, and Dovitinib) were added to the cell cultures in a volume of 300μl (for PC3) and 1 ml (for DU145 and LNCaP) to maintain the constant volume. Experiments were performed in triplicate.

From the initial plating of single cells iPS87 at 8.0 x 10^4^ cells/ml using 12-well TC plates, measurements were taken on days 1, 4, 7, and 11 and the inhibitors (BGJ398, and Dovitinib) were added to the cell cultures every 3 days (on days 1, 4, and 7). Experiments were performed in biological triplicates with technical duplicate samples.

### RT-qPCR reagents and primers

Cells were collected and washed with chilled 1x PBS and then RNA was extracted using Trizol reagent (Invitrogen) according to the manufacturer’s protocol. Total RNA concentration was measured using nanodrop. 100 ng of total RNA was used to prepare cDNA using ProtoScript II Reverse Transcriptase (NEB #M0368) with oligo(dT) primers (Cat# 51-01-15-01, IDT) following the manufacturer’s protocol. qPCR was performed using a SYBR green assay system with Phusion® High-Fidelity DNA Polymerase using a Stratagene Mx3000 qPCR machine (Agilent Technologies, CA, USA). The mRNA levels were normalized to beta-actin abundance, and the fold change between samples was calculated by a standard ΔΔCt analysis.

Following primers were used: FGFR1 forward 5’-TAATGGACTCTGTGGTGCCCTC-3’, reverse 5’-ATGTGTGGTTGATGCTGCCG-3’ ^45^; OCT4 forward 5’-GCAATTTGCCAAGCTCCTGAA-3’, reverse 5’-GCAGATGGTCGTTTGGCTGA-3’ ^50^; ALDH7A1 forward 5′-CAACGAGCCAATAGCAAGAG-3′, reverse 5′-GCATCGCCAATCTGTCTTAC-3′ ^10^; E-cadherin forward 5′-CGGGAATGCAGTTGAGGATC-3′, reverse 5′-AGGATGGTGTAAGCGATGGC-3′ ^51^; vimentin forward 5′-AGATGGCCCTTGACATTGAG -3′, reverse 5′-TGGAAGAGGCAGAGAAATCC-3′ ^52^; Snail forward 5′-GAAAGGCCTTCAACTGCAAA-3′, reverse 5′-TGACATCTGAGTGGGTCTGG-3′ ^51^; beta-actin forward 5’-AGAGCTACGAGCTGCCTGAC-3’, reverse 5’-AGCACTGTGTTGGCGTACAG-3’ ^53^.

### Immunoblotting, antibodies and additional reagents

Lysates were collected in radioimmunoprecipitation assay buffer [RIPA; 50 mM Tris-HCl (pH 8.0), 150 mM NaCl, 1% TritionX-100, 0.5% sodium deoxycholate, 0.1% SDS, 50 mM NaF, 1 mM sodium orthovanadate, 1 mM PMSF, and 10 μg/mL aprotinin]. 25 μg or 30 μg of total protein was separated by 10% or 12.5% SDS-PAGE followed by transfer to Immobilon-P membrane. Immunoblotting reagents were from the following sources: antibodies against p-FGFR (Tyr653/654), FGFR1 (D8E4), FGFR4 (D3B12), p-VEGFR (19A10), VEGFR2 (55B11), p-AKT (D9E), AKT (9272), p-MAPK (D13.14.4E), p44/42 MAPK (Erk1/2), OCT4 (2750), CD133 (D2V8Q), Androgen Receptor (D6F11), and β-actin (4967) antibodies were from Cell Signaling Technology; FGFR2 (C-8), FGFR3 (B-9), and STAT5 (C-17) were from Santa Cruz Biotechnology; ALDH7A1 (CAT:ABO11656) was from Abgent; HRP anti-mouse, HRP anti-rabbit, and Enhanced Chemiluminescence (ECL) reagents were from GE Healthcare. Other reagents included: Dovitinib, BGJ398 and PD166866 were from Selleckchem.

## ACKNOWLEDGEMENTS

We thank all current lab members particularly Nicole Peiris for encouragement and invaluable editing; Christy Cho and Prof. Neal Devraj at UCSD for microscopic cell imaging; and Vince Harjono and Prof. Brian Zid at UCSD for help performing qPCR experiments. DJD gratefully acknowledges generous philanthropic support from the UC San Diego Foundation.

## REFERENCES

1. Bray F, Ferlay J, Soerjomataram I, Siegel RL, Torre LA, Jemal A. Global cancer statistics 2018: GLOBOCAN estimates of incidence and mortality worldwide for 36 cancers in 185 countries. CA Cancer J Clin. 2018;68(6):394–424.

2. Smith MR, Saad F, Chowdhury S, et al. Apalutamide Treatment and Metastasis-free Survival in Prostate Cancer. N Engl J Med. 2018;378(15):1408–1418.

3. Urbanucci A, Sahu B, Seppala J, et al. Overexpression of androgen receptor enhances the binding of the receptor to the chromatin in prostate cancer. Oncogene. 2012;31(17):2153–2163.

4. Fennema E, Rivron N, Rouwkema J, van Blitterswijk C, de Boer J. Spheroid culture as a tool for creating 3D complex tissues. Trends Biotechnol. 2013;31(2):108–115.

5. Kim JB. Three-dimensional tissue culture models in cancer biology. Semin Cancer Biol. 2005;15(5):365–377.

6. Portillo-Lara R, Alvarez MM. Enrichment of the Cancer Stem Phenotype in Sphere Cultures of Prostate Cancer Cell Lines Occurs through Activation of Developmental Pathways Mediated by the Transcriptional Regulator DeltaNp63alpha. PLoS One. 2015;10(6):e0130118.

7. Liao WT, Ye YP, Deng YJ, Bian XW, Ding YQ. Metastatic cancer stem cells: from the concept to therapeutics. Am J Stem Cells. 2014;3(2):46–62.

8. Bahmad HF, Cheaito K, Chalhoub RM, et al. Sphere-Formation Assay: Three-Dimensional in vitro Culturing of Prostate Cancer Stem/Progenitor Sphere-Forming Cells. Front Oncol. 2018;8:347.

9. Reya T, Morrison SJ, Clarke MF, Weissman IL. Stem cells, cancer, and cancer stem cells. Nature. 2001;414(6859):105–111.

10. van den Hoogen C, van der Horst G, Cheung H, Buijs JT, Pelger RC, van der Pluijm G. The aldehyde dehydrogenase enzyme 7A1 is functionally involved in prostate cancer bone metastasis. Clin Exp Metastasis. 2011;28(7):615–625.

11. Kapalczynska M, Kolenda T, Przybyla W, et al. 2D and 3D cell cultures - a comparison of different types of cancer cell cultures. Arch Med Sci. 2018;14(4):910–919.

12. Edmondson R, Broglie JJ, Adcock AF, Yang L. Three-dimensional cell culture systems and their applications in drug discovery and cell-based biosensors. Assay Drug Dev Technol. 2014;12(4):207–218.

13. Hirschhaeuser F, Menne H, Dittfeld C, West J, Mueller-Klieser W, Kunz-Schughart LA. Multicellular tumor spheroids: an underestimated tool is catching up again. J Biotechnol. 2010;148(1):3–15.

14. Ornitz DM, Itoh N. The Fibroblast Growth Factor signaling pathway. Wiley Interdiscip Rev Dev Biol. 2015;4(3):215–266.

15. Porta R, Borea R, Coelho A, et al. FGFR a promising druggable target in cancer: Molecular biology and new drugs. Crit Rev Oncol Hematol. 2017;113:256–267.

16. Corn PG, Wang F, McKeehan WL, Navone N. Targeting fibroblast growth factor pathways in prostate cancer. Clin Cancer Res. 2013;19(21):5856–5866.

17. Gallo LH, Nelson KN, Meyer AN, Donoghue DJ. Functions of Fibroblast Growth Factor Receptors in cancer defined by novel translocations and mutations. Cytokine Growth Factor Rev. 2015;26(4):425–449.

18. Li B, Sun A, Youn H, et al. Conditional Akt activation promotes androgen-independent progression of prostate cancer. Carcinogenesis. 2007;28(3):572–583.

19. Cassinelli G, Zuco V, Gatti L, et al. Targeting the Akt kinase to modulate survival, invasiveness and drug resistance of cancer cells. Current medicinal chemistry. 2013;20(15):1923–1945.

20. Qian X, Anzovino A, Kim S, et al. N-cadherin/FGFR promotes metastasis through epithelial-to-mesenchymal transition and stem/progenitor cell-like properties. Oncogene. 2014;33(26):3411–3421.

21. Zhao Q, Parris AB, Howard EW, et al. FGFR inhibitor, AZD4547, impedes the stemness of mammary epithelial cells in the premalignant tissues of MMTV-ErbB2 transgenic mice. Scientific reports. 2017;7(1):11306.

22. Vasiutkov V. [Damage to the major vessels during surgical interventions in patients with cancer]. Vopr Onkol. 1988;34(12):1485–1489.

23. Maehara O, Suda G, Natsuizaka M, et al. Fibroblast growth factor-2-mediated FGFR/Erk signaling supports maintenance of cancer stem-like cells in esophageal squamous cell carcinoma. Carcinogenesis. 2017;38(11):1073–1083.

24. Armstrong K, Ahmad I, Kalna G, et al. Upregulated FGFR1 expression is associated with the transition of hormone-naive to castrate-resistant prostate cancer. Br J Cancer. 2011;105(9):1362–1369.

25. Sahadevan K, Darby S, Leung HY, Mathers ME, Robson CN, Gnanapragasam VJ. Selective over-expression of fibroblast growth factor receptors 1 and 4 in clinical prostate cancer. J Pathol. 2007;213(1):82–90.

26. Acevedo VD, Gangula RD, Freeman KW, et al. Inducible FGFR-1 activation leads to irreversible prostate adenocarcinoma and an epithelial-to-mesenchymal transition. Cancer Cell. 2007;12(6):559–571.

27. Wan X, Corn PG, Yang J, et al. Prostate cancer cell-stromal cell crosstalk via FGFR1 mediates antitumor activity of dovitinib in bone metastases. Sci Transl Med. 2014;6(252):252ra122.

28. Finones RR, Yeargin J, Lee M, et al. Early human prostate adenocarcinomas harbor androgen-independent cancer cells. PLoS One. 2013;8(9):e74438.

29. Assoun EN, Meyer AN, Jiang MY, Baird SM, Haas M, Donoghue DJ. Characterization of iPS87, a prostate cancer stem cell-like cell line. Oncotarget. 2020;11(12):1075–1084.

30. Yu SQ, Lai KP, Xia SJ, Chang HC, Chang C, Yeh S. The diverse and contrasting effects of using human prostate cancer cell lines to study androgen receptor roles in prostate cancer. Asian J Androl. 2009;11(1):39–48.

31. Feng S, Shao L, Yu W, Gavine P, Ittmann M. Targeting fibroblast growth factor receptor signaling inhibits prostate cancer progression. Clin Cancer Res. 2012;18(14):3880–3888.

32. Mori S, Chang JT, Andrechek ER, et al. Anchorage-independent cell growth signature identifies tumors with metastatic potential. Oncogene. 2009;28(31):2796–2805.

33. Shi X, Gipp J, Bushman W. Anchorage-independent culture maintains prostate stem cells. Dev Biol. 2007;312(1):396–406.

34. Wagner K, Welch D. Cryopreserving and recovering of human iPS cells using complete Knockout Serum Replacement feeder-free medium. J Vis Exp. 2010(41).

35. Roberts E, Cossigny DA, Quan GM. The role of vascular endothelial growth factor in metastatic prostate cancer to the skeleton. Prostate Cancer. 2013;2013:418340.

36. Lee JT, Steelman LS, Chappell WH, McCubrey JA. Akt inactivates ERK causing decreased response to chemotherapeutic drugs in advanced CaP cells. Cell Cycle. 2008;7(5):631–636.

37. Brocker C, Lassen N, Estey T, et al. Aldehyde dehydrogenase 7A1 (ALDH7A1) is a novel enzyme involved in cellular defense against hyperosmotic stress. J Biol Chem. 2010;285(24):18452–18463.

38. Rizzino A, Wuebben EL. Sox2/Oct4: A delicately balanced partnership in pluripotent stem cells and embryogenesis. Biochim Biophys Acta. 2016.

39. Zeineddine D, Hammoud AA, Mortada M, Boeuf H. The Oct4 protein: more than a magic stemness marker. Am J Stem Cells. 2014;3(2):74–82.

40. van den Hoogen C, van der Horst G, Cheung H, et al. High aldehyde dehydrogenase activity identifies tumor-initiating and metastasis-initiating cells in human prostate cancer. Cancer Res. 2010;70(12):5163–5173.

41. Panek RL, Lu GH, Dahring TK, et al. In vitro biological characterization and antiangiogenic effects of PD 166866, a selective inhibitor of the FGF-1 receptor tyrosine kinase. J Pharmacol Exp Ther. 1998;286(1):569–577.

42. Yadav SS, Li J, Stockert JA, et al. Induction of Neuroendocrine Differentiation in Prostate Cancer Cells by Dovitinib (TKI-258) and its Therapeutic Implications. Transl Oncol. 2017;10(3):357–366.

43. Strouhalova K, Prechova M, Gandalovicova A, Brabek J, Gregor M, Rosel D. Vimentin Intermediate Filaments as Potential Target for Cancer Treatment. Cancers (Basel). 2020;12(1).

44. Jiang MY, Lee TL, Hao SS, et al. Visualization of early prostatic adenocarcinoma as a stem cell disease. Oncotarget. 2016;7(46):76159–76168.

45. Ji W, Yu Y, Li Z, et al. FGFR1 promotes the stem cell-like phenotype of FGFR1-amplified non-small cell lung cancer cells through the Hedgehog pathway. Oncotarget. 2016;7(12):15118–15134.

46. Bluemn EG, Coleman IM, Lucas JM, et al. Androgen Receptor Pathway-Independent Prostate Cancer Is Sustained through FGF Signaling. Cancer Cell. 2017;32(4):474–489 e476.

47. De Marzo AM, Knudsen B, Chan-Tack K, Epstein JI. E-cadherin expression as a marker of tumor aggressiveness in routinely processed radical prostatectomy specimens. Urology. 1999;53(4):707–713.

48. Bae KM, Su Z, Frye C, et al. Expression of pluripotent stem cell reprogramming factors by prostate tumor initiating cells. J Urol. 2010;183(5):2045–2053.

49. Hepburn AC, Curry EL, Moad M, et al. Propagation of human prostate tissue from induced pluripotent stem cells. Stem Cells Transl Med. 2020;9(7):734–745.

50. Bae WJ, Lee SH, Rho YS, Koo BS, Lim YC. Transforming growth factor beta1 enhances stemness of head and neck squamous cell carcinoma cells through activation of Wnt signaling. Oncol Lett. 2016;12(6):5315–5320.

51. Wang W, Wang L, Mizokami A, et al. Down-regulation of E-cadherin enhances prostate cancer chemoresistance via Notch signaling. Chin J Cancer. 2017;36(1):35.

52. Avtanski D, Garcia A, Caraballo B, et al. In vitro effects of resistin on epithelial to mesenchymal transition (EMT) in MCF-7 and MDA-MB-231 breast cancer cells - qRT-PCR and Westen blot analyses data. Data Brief. 2019;25:104118.

53. Chen F, Zhang G, Yu L, et al. High-efficiency generation of induced pluripotent mesenchymal stem cells from human dermal fibroblasts using recombinant proteins. Stem Cell Res Ther. 2016;7(1):99.

